# The Role of Ribose Modifications on the Structural Stability of Nucleotide Analogs with α-thiotriphosphate at the Active Site of SARS-CoV-2 RdRp

**DOI:** 10.1101/2025.04.11.648322

**Authors:** Yongfang Li, Shiling Yuan

## Abstract

As a promising drug target, RNA-dependent RNA polymerase (RdRp) has attracted much attention recently due to its notable conserved active site, especially in the context of COVID-19 spreading. To inhibit the function of RdRp, nucleotide analog is a common choice for acting as a chain terminator or RNA corruptor. Although some nucleotide analogs have shown the ability to terminate the extension of a nascent strand of SARS-CoV-2, most of them are likely to be excised due to the proofreading of SARS-CoV-2 nsp14/nsp10. A previous experimental study found that introducing sulfur modification into analogs’ phosphate moieties (α-thiotriphosphate; “thio” modification) can break the bottleneck. For instance, Sofosbuvir with α-thiotriphosphate modification successfully escaped from the excision of nsp14/10. However, it is unknown how the α-thiotriphosphate affects the structural stability of nucleotide analogs with different ribose modifications at the active site of SARS-CoV-2 RdRp. Thus, in this study, we performed extensive molecular dynamics simulations on four nucleotide analogs with α-thiotriphosphate to elucidate what kind of ribose modification combined with “thio” modification would benefit the analog’s structural stability at the active site. We found that the “thio” modification led to the torsion of phosphate moieties, profoundly affecting the overall conformation of analogs and surrounding residues, determining the Watson-Crick base pairing and catalytic efficiency. Interestingly, chemical modification on the ribose, especially the 1’ and 3’-ribose positions, increases the structural stability of analogs through hydrogen bond interactions. Our results revealed nucleotide analogs’ structural and dynamical features with “thio” modification at the active site, which may contribute to future drug design or repurposing aimed at the SARS-CoV-2 RdRp.

## Introduction

SARS-CoV-2 caused a global pandemic from 2019 to 2023, leading to more than 676 million infection cases and over 6.8 million deaths in 196 countries^1^. So far, there are no fully effective drugs to combat SARS-CoV-2 and its variants efficiently^2, 3^. Thus, the development of antiviral drugs is still necessary for the treatment of infected cases. A promising way to combat the virus is screening or re-designing the antiviral drugs from the current drug database instead of developing new drugs from scratch, which could save time and cost^4-8^. The nucleoside/tide analogs (NA) have already shown a high potential to inhibit virus replication^9-12^.

NA is a class of compounds with chemical modification on the ribose ring, base ring, or both compared with natural nucleosides/tide^4, 9, 13^. In combating coronavirus, NA mainly aims to inhibit the function of RNA-dependent RNA polymerase (RdRp), an essential enzyme for the virus’s life cycle^14-18^. RdRp relies on the nucleoside addition cycle to carry on the replication or transcription of the virus gene^19^. NA can disturb the nucleoside addition cycle of SARS-CoV-2 RdRp by loading into the nascent strand RNA, acting as an inhibitor or corruptor^20^. In the current form of NAs, there are usually three positions of ribose that can be modified to other chemical groups, named 1’, 2’, and 3’ positions. For instance, a famous 1’ modified NAs is Remdesivir, which has a cyano modification at the 1’ position of ribose^21-23^. It is used to treat outpatients at the early stage of infection and patients hospitalized due to the COVID-19^24-26^, approved by the U.S. Food and Drug Administration (FDA). A common modification strategy for NA is replacing or removing the hydroxyl group on the ribose’s 2’ or 3’ position. An experimental study in vitro carried out by Ju et al. demonstrated that Sofosbuvir in triphosphate form, a nucleotide analog with 2’-ribose modification, can terminate the RNA synthesis catalyzed by the SARS-CoV-2 RdRp^10^. Further simulation research revealed that incorporating Sofosbuvir into the nascent RNA strand led to immediate chain termination due to the apparent steric effect introduced by the bulky 2’-methyl groups on the ribose^27^. Another simulation research investigated two similar NAs in adenosine form, Didanosine and Stavudine, which presented feasible structural stability at the active site of SARS-CoV-2 RdRp^28^. When incorporated into the nascent RNA strand, analogs can inhibit the incorporation of the following natural NTP. They proposed that an appropriate combination of modification, such as polar group on 2’-ribose and blocking the 3’-ribose, would offer the analogs the ability of immediate termination on the nascent RNA chain.

However, reasonable modifications on the ribose or base do not guarantee that the NA effectively inhibits the SARS-CoV-2 replication^29^. Although some NA present inhibition on the elongation of SARS-CoV-2 RdRp, NA can be excised by the 3’-to-5’ exoribonuclease (ExoN) in nonstructural protein 14 (nsp14)^30-32^. This coronavirus proofreading process guarantees the RNA genome’s high-fidelity replication^33, 34^. Recent reports demonstrated that NA without the 3’-hydroxyl group on the ribose has a limited effect on inhibiting the replication of SARS-CoV-2^20, 35^. Crystal structure analysis proposed that the 2’ and 3’-hydroxyl groups on the ribose may be responsible for the structural stability of NTP/analogs at the active site of RdRp^19, 36, 37^. It is unknown if lacking the 3’-hydroyxl group of analogs at the active site could trigger the excision of ExoN. Besides, even with the presence of the 3’-hydroxyl group, the inhibition effect of NA is limited. For example, although sofosbuvir can terminate the extension of the nascent chain synthesized by the RdRp, the ability to terminate the chain extension disappeared when ExoN existed^30^.

Thus, nucleoside analogs aimed at SARS-CoV-2 RdRp have to resist the proofreading of ExoN. To this end, previous studies have attempted to combine NA with other drugs that can inhibit the activity of ExoN to terminate the elongation of RdRp. Experimental research in vitro carried out by Souza et al. demonstrated that two drugs, Pibrentasvir and Ombitasvir, inhibit the SARS-CoV-2 exonuclease in a concentration-dependent manner^38^. Compared with the absence of Pibrentasivir, multiple NA, including Sofosbuvir and Molnupiravir, have a lower chance of being excised by exonuclease with the presence of Pibrentasivir. Moreover, a clinically artificial intelligence platform combined with experimental validation of multi-drug efficacy on a SARS-CoV-2 proposed that Ritonavir and Lopinavir with Remdesivir may be effective drug combinations aimed at the SARS-CoV-2 RdRp and ExoN^39^. Nevertheless, careful dose adjustment and optimization strategies are necessary for further preclinical and/or clinical evaluation to maximize synergistic efficacy at minimal toxicity.

Although drug combinations therapy shade light on the drug screening aimed at SARS-CoV-2 RdRp and ExoN, those NA directly inhibiting the activity of SARS-CoV-2 ExoN may provide higher efficiency on the inhibition of SARS-CoV-2 RdRp. Recently, Canard et al. synthesized a new NA named AT-9052, the α-thiotriphosphate form of 5’-TP AT-9010^40^. As a potent chain terminator, AT-9010 is easily excised by the ExoN due to the backtracking or stalling of replication/transcription complex (RTC), triggering the association of nsp14^41^. The compound in the form of α-thiotriphosphate (AT-9052) can be incorporated into the nascent strand as substrate, which would terminate the chain extension. Meanwhile, the *S*_*P*_ form of AT-9052 shows resistance to the excision by the SARS-CoV-2 nsp14/nsp10. Based on the rich combination of modification on the 2’ and 3’-ribose of NA, the form of α-thiotriphosphate may unlock the considerable potential of NA, whose activity is limited by the proofreading of ExoN. More importantly, this method allows a single NA to inhibit RdRp and ExoN activities.

It is unknown what new impaction of the “thio” modification brings to the NA with different ribose modifications at the active site of SARS-CoV-2 RdRp. Although the experimental results demonstrated that “thio” modification on the NA may lower the reaction rate of NA incorporating into the nascent chain, more NA with diverse ribose or base modification are supposed to be tested to uncover what kind of ribose modification combined with “thio” modification would benefit the NA incorporation efficiency at the SARS-CoV-2 RdRp^40^. Investigating NA’s structural stability and dynamic features in the α-thiotriphosphate at the RdRp active site is necessary. To this end, four purine nucleoside analogs, including Cladribine, Cordycepin, 2’-NH_2_-ATP, and Remdesivir with “thio” modification, were investigated in this study. The abbreviations for the four nucleoside analogs are CAS, CRS, NHS, and RES (Figure 1).

**Figure 1.**
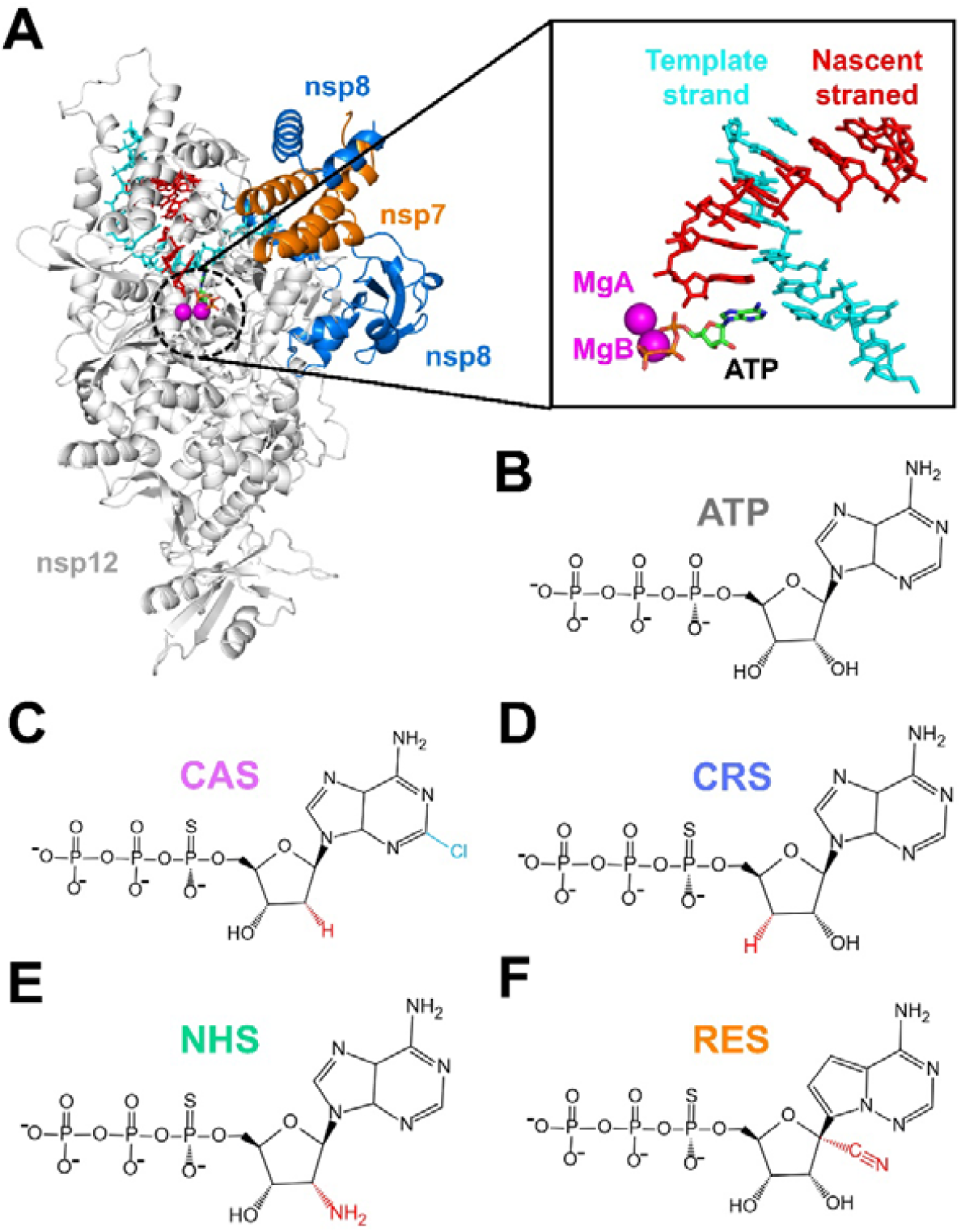
(A) Diagram of SARS-CoV-2 RdRp, where ATP binding at the active site is enlarged. (B–F) Compared to adenosine in triphosphate form, the chemical structures of four nucleotide analogs in α-thiotriphosphate form, where ribose modifications are highlighted in red and base modifications in blue.

Cladribine was approved for treating hairy cell leukemia with chlorine modification on the base ring, and the hydroxyl group on the 2’-ribose position was canceled off^42^. Cordycepin canceled off the hydroxyl group on the 3’-ribose position, a potential “immediate” inhibitor of the SARS-CoV-2 RdRp proposed by previous experimental and computational studies^28, 43^. 2’-NH_2_-ATP has an amino group on the 2’-ribose position, which provides potential hydrogen bond interaction with surrounding residues^9, 28^. Remdesivir was approved for treating various RNA virus infections, including COVID-19, and the 1’-ribose position was modified to the cyano group^44^. Distance distribution, hydrogen bond analysis, angle distribution, and root mean squared fluctuation (RMSF) were applied to elucidate how the “thio” modification affects the structural stability of analogs at the active site and the role of ribose/base modification of analogs. We found that sulfur modification of analogs would cause obvious torsion on the phosphate moieties, leading to a change in catalytic conformations and the disruption of Watson-Crick base pairing. Meanwhile, chemical modifications on the 1’ and 3’-ribose positions play a vital role in the structural stability of analogs at the active site. Our discovery provides a detailed dynamic picture of nucleotide analogs with “thio” modification at the SARS-CoV-2 RdRp, demonstrating that “thio” modification combined with various ribose modifications of analogs may be a hopeful way to inhibit the SARS-CoV-2.

## Methods

The simulation systems were built from the crystal structure of SARS-CoV-2 RdRp (PDB ID: 7AAP), in which two Mg^2+^ ions coordinated at active sites and in complex with favipiravir in triphosphate form^45^. The favipiravir at the active site was mutated to ATP and other analogs in the α-thiotriphosphate form, and the corresponding base pairing was mutated to the uracil. The missing residues were filled by the Modeller 9.21^46^. The PROPKA web server predicted the protonation state of the complex^47^. The interaction between the Zn^2+^ ions and corresponding histidine residues was coordinated by manual inspection. Each complex was placed into a dodecahedron box with the box edges at least 12 Å away from the complex surface. The TIP3P water model was used to solvate the complex^48^, and sufficient ions were added to neutralize the whole system.

The Amber FF14SB force field with RNA_OL3 corrections simulated the protein and nucleic^49-51^. The force field parameter of ATP applied in this paper followed the previous research^52, 53^. The parameters of four nucleotide analogs were generated using the General Amber Force Field (GAFF)^54^. The structures of nucleotide analogs were optimized at the Hartree–Fock (HF) level and the 6-31G(d,p) basis set, and then their partial charges were determined by the RESP method using Antechamber tools^55, 56^. Structural optimization and energy calculation at the quantum chemistry level were performed using the Gaussian16 package^57^.

For each simulation system, two rounds of 10,000 steps of energy minimization were performed with and without position restraints on the heavy atoms of dsRNA, respectively. We then performed 1 ns NPT equilibration simulation (T=300 K; P=1 bar) with position restraints on all the heavy atoms with a force constant of 10 kJ×mol^-1^ ×Å^-2^. Afterward, we removed the position restraints and harmonic constraints for Mg^2+^ in the active site. The last configuration of the NPT equilibration simulation was used to perform a 50 ns NPT (T=300 K; P=1 bar) simulation followed by a 100 ns NVT (T=300 K) simulation to reach the equilibrium. The last conformation of NVT simulation was used for the seed MD production simulations under the NVT ensemble (T=300K). For each system, 10×50ns trajectories were generated, and the snapshot was saved every 20 ps. The integration time step was 2 fs, and the neighbor list was updated every 10 steps. The cut-off of the Van der Waals interactions is 12Å. The short-range electrostatic interactions were cut off at 12Å, and the Particle-Mesh Ewald method was used to treat the long-range electrostatic interaction^58^. All bonds were constrained using LINC algorithms^59^. The V-rescale thermostat was used to control the temperature at 300 K with a coupling time of 0.1 ps^60^. All the MD simulations were performed by GROMACS 2022.03^61^.

In this work, the bootstrap algorithm was adopted to calculate the mean value and the standard deviations, including Hbond probability and RMSF. Specifically, we generated 10 bootstrap samples for each system for the calculation. For each bootstrap sample, 10 trajectories were chosen randomly from the conformational ensemble.

## Results and Discussion

To uncover the impact of “thio” modification on the analogs, we first investigated the conformational changes of analogs at the active site of SARS-CoV-2 RdRp by angle distribution (Figure 2). The dihedral angle distribution is defined by atoms on the analogs and Mg^2+^ ions or 3’-terminal RNA, providing an overall picture of the analog structural stability at the active site. Due to different chemical modifications, each analog was divided into three areas to carry on the analysis, including phosphate moieties, ribose ring, and base ring. The non-bridge oxygen atom on the P_α_ was replaced by a sulfur atom in the “thio” modification.

**Figure 2.**
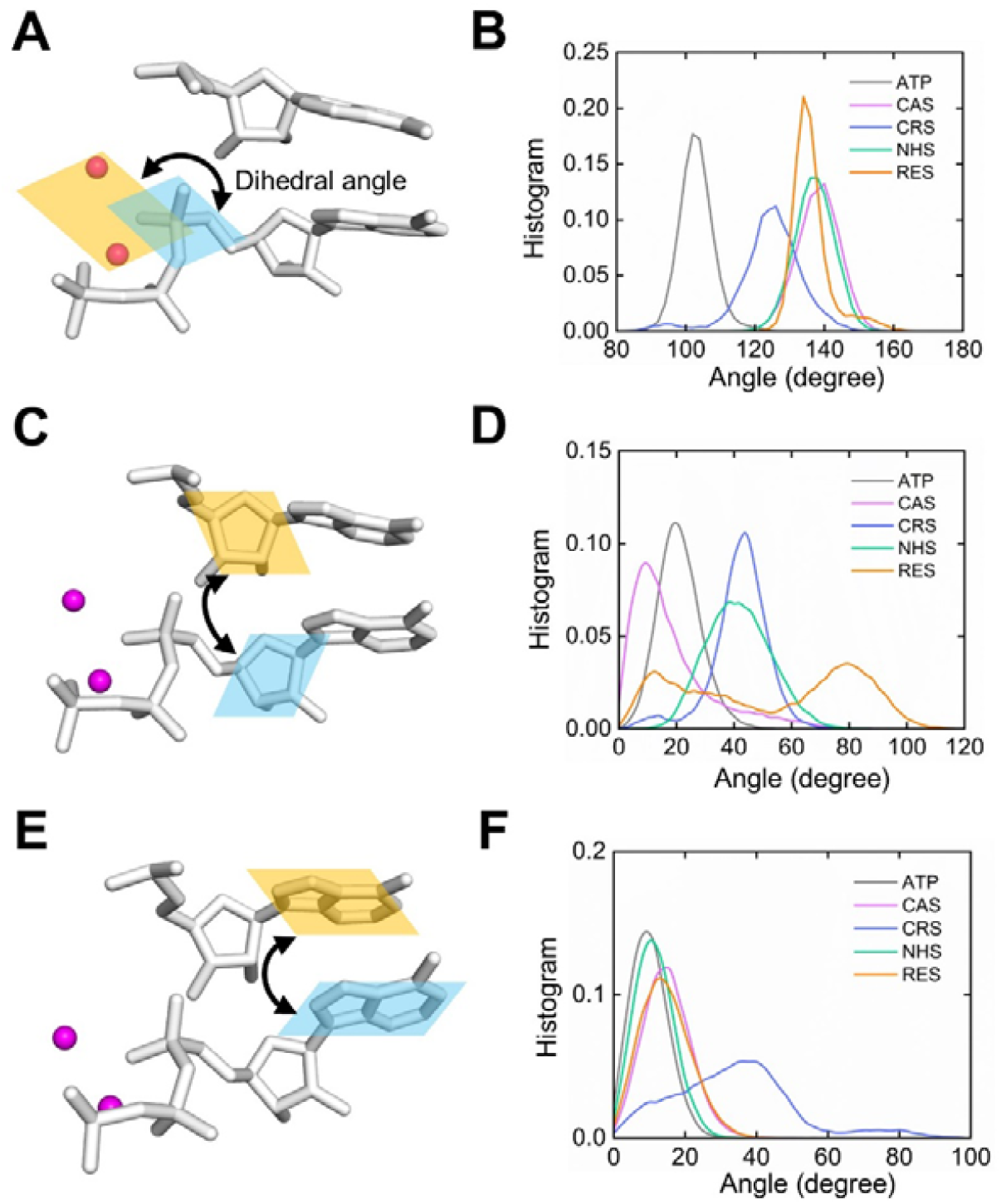
Examinations of the dihedral angle of different parts of nucleotide analogs between the active site (*i* site) and *i*+1 site on the nascent strand. (A) Diagram showing the two planes used to calculate the dihedral angle. The first plane is defined by two Mg^2+^ ions and P_α_ atom. The second plane is defined by three atoms connected to the P_α_ atom. (B) Angle distribution of phosphate moieties for different analogs. (C) Diagram of ribose angle between *i* site and *i*+1 site. The ribose plane was defined by atoms on the ribose’s 1’, 4’, and 5’ positions. (D) Angle distribution of ribose ring for different analogs. (E) Diagram of the base angle between *i* site and *i*+1 site. (F) Angle distribution of base ring for different analogs.

First, we check the orientation of the phosphate moieties related to the Mg^2+^ ions (Figure 2A). Two Mg^2+^ ions and P_α_ atoms define the first plane; the second is defined by three atoms bonded to P_α_. Compared to the natural ATP system, the angle distribution of four analogs presents an evident shift, concentrating at 135 degrees. Then, we checked the distribution of the ribose angle defined by the ribose at the active site (*i* site) and that of the *i*+1 site. Analog systems show obvious shifted distributions from the natural ATP system except for the CAS system. We noticed that the RES system presents two distribution peaks, and one of the peaks overlapped with that of the ATP system. We also checked the distribution of the base angle defined by the base ring at the active site and *i*+1 site. Three analog systems, CAS, NHS, and RES, show the same distribution as the ATP system, while CRS presents different distributions from the ATP system.

Based on the above angle distribution, we observed that different chemical modifications of analogs offer diverse conformational distribution at the SARS-CoV-2 RdRp active site when they are in the “thio” modification. Those conformational changes related to the catalytical activities play an essential role in the inhibition efficiency of analogs^28, 62^. It is already known that the sulfur atom bonded to the P_α_ introduced torsion of phosphate moieties to the analogs. To uncover the detailed pictures of the catalytical conformation of different analogs at the active site, we calculated multiple distance distributions defined by Mg^2+^ ions and nucleic atoms (Figure 3), including the distance of P_α_-O3’, MgA-P_α_, and MgA-O3’. For the distance distribution of Pα-O3’, all four analogs show a distribution of greater distance than the natural ATP system. RES compound presents two distribution peaks, one located in 4.5 Å overlapped with the NHS compound. The distribution peak of CAS is 4 Å, which is slightly closer to the peak of the ATP system. CRS system shows the greatest distance with the peak position at 8 Å. The MgA-Pα distance of four analogs is greater than that of ATP. The peak positions of CAS, CRS, and NHS are beyond 5 Å, while that of RES compound is less than 4 Å. The distances of MgA-O3’ in the analog systems are also affected by the sulfur modification, which is greater than 4 Å, with only a few conformations of RES aligned with the natural ATP system.

**Figure 3.**
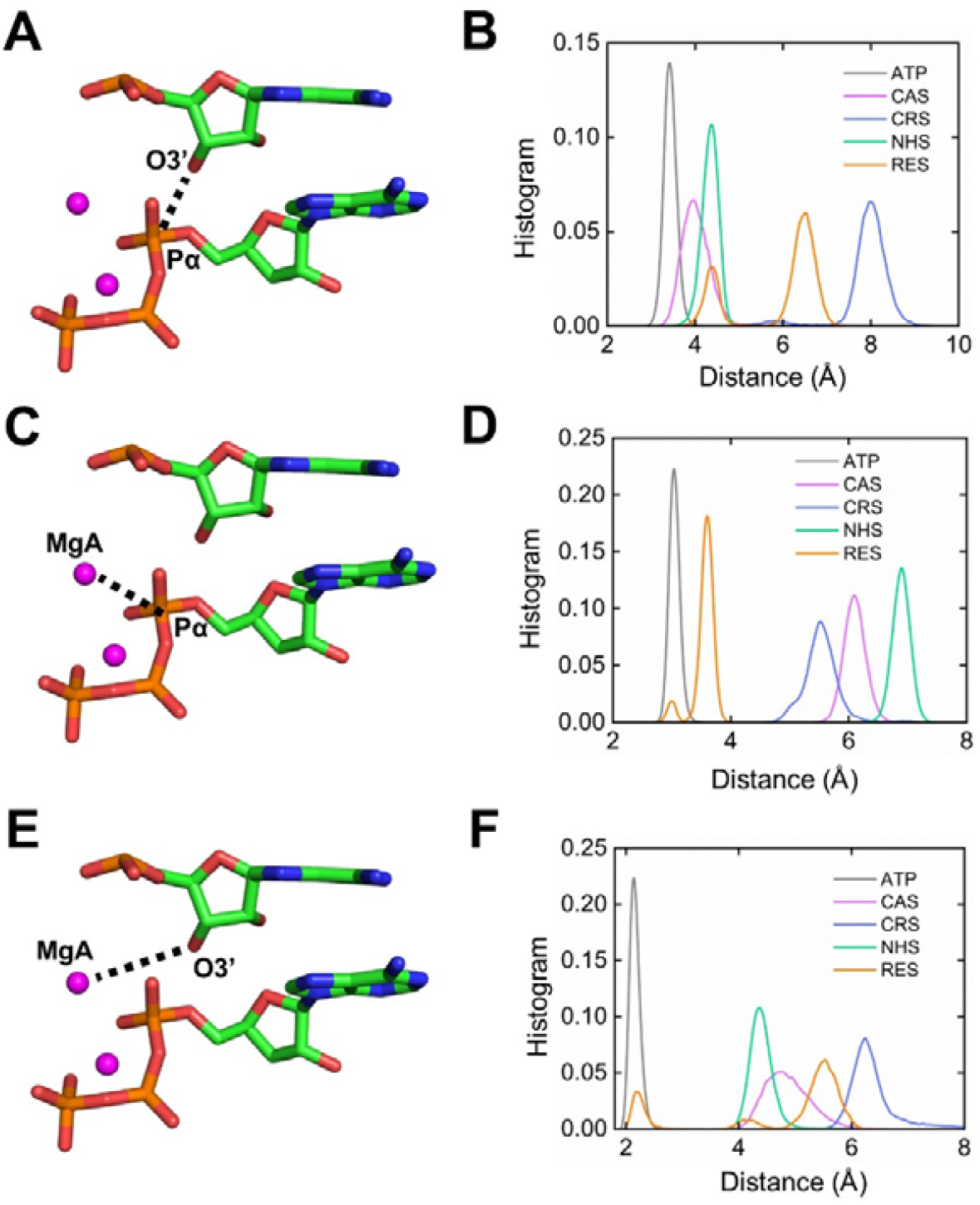
Investigation of the distance distribution on the catalytic conformations for different nucleotide analogs. (A) Diagram showing the distance between the P_α_ atom of analogs and the O3’ atom of the 3’-terminal of the nascent strand. (B) Distance distribution of P_α_-O3’ for analog systems compared with ATP system. (C) Diagram showing the distance between the P_α_ atom of analogs and the MgA ion at the active site. (D) Distance distribution of P_α_-MgA for analogs systems compared with ATP system. (E) Diagram showing the distance between the MgA ion at the active site and the O3’ atom of the 3’-terminal of the nascent strand. (F) Distance distribution of MgA-O3’ for analogs systems compared with ATP system.

A previous study pointed out that the distance Pα-O3’ is required to be less than 4 Å for efficient phosphodiester bond formation during the catalysis reaction^63^. Compared with natural nucleotide, all four analogs with sulfur modification on the phosphate moieties do not possess complete catalytical ability. This is aligned with the experimental results that analogs in “thio” modification showed a 2.8-fold decrease in the catalytic rate compared to analogs with triphosphate moieties^40^. Moreover, the distribution of MgA-Pα, and MgA-O3’ demonstrated that the coordination of Mg^2+^ ions was disturbed by the torsion of analogs phosphate moieties. We also checked the distance between two Mg^2+^ ions and found that this distance of all analog systems is greater than that of the ATP system (Figure S1). The above distance results imply that the catalytic conformation of analogs in “thio” modification at the active site is looser than that of natural nucleotide with normal triphosphate. This may lead to unexpected conformational changes in surrounding residues.

Previous studies pointed out that Arg555 is a highly conserved flexible residue adopting a wide range of positions at the RdRp active site^14, 37, 64^. It recognizes incoming NTP by interacting with the NTP base or β-phosphate oxygens^37^. Thus, we investigated the interaction distance from the side chain of Arg555 to the base of analogs and the β-phosphate oxygens (O_Pβ_), respectively (Figure 4). For the natural ATP system, the interaction of Arg555-O_Pβ_ is relatively stable, with over 95.4±2.4% conformations having a distance below 4 Å, while the distribution of Arg555-Base is wide. RES system shows a similar distance distribution to the ATP system, except that the Arg555-O_Pβ_ distribution is more concentrated. CAS system shows a relatively large distribution for both distances, demonstrating that the Arg555 is flexible during the simulation. For the CRS system, the Arg555-Base distance is concentrated on the 3.5 Å, while the Arg555-O_Pβ_ interaction is very weak, with most of the conformations having a distance beyond 6 Å. The interaction of Arg555-Base is stronger than that of Arg555-O_Pβ_ in the NHS system, with distance concentrated on the 3.5 Å and 4 Å, respectively.

**Figure 4.**
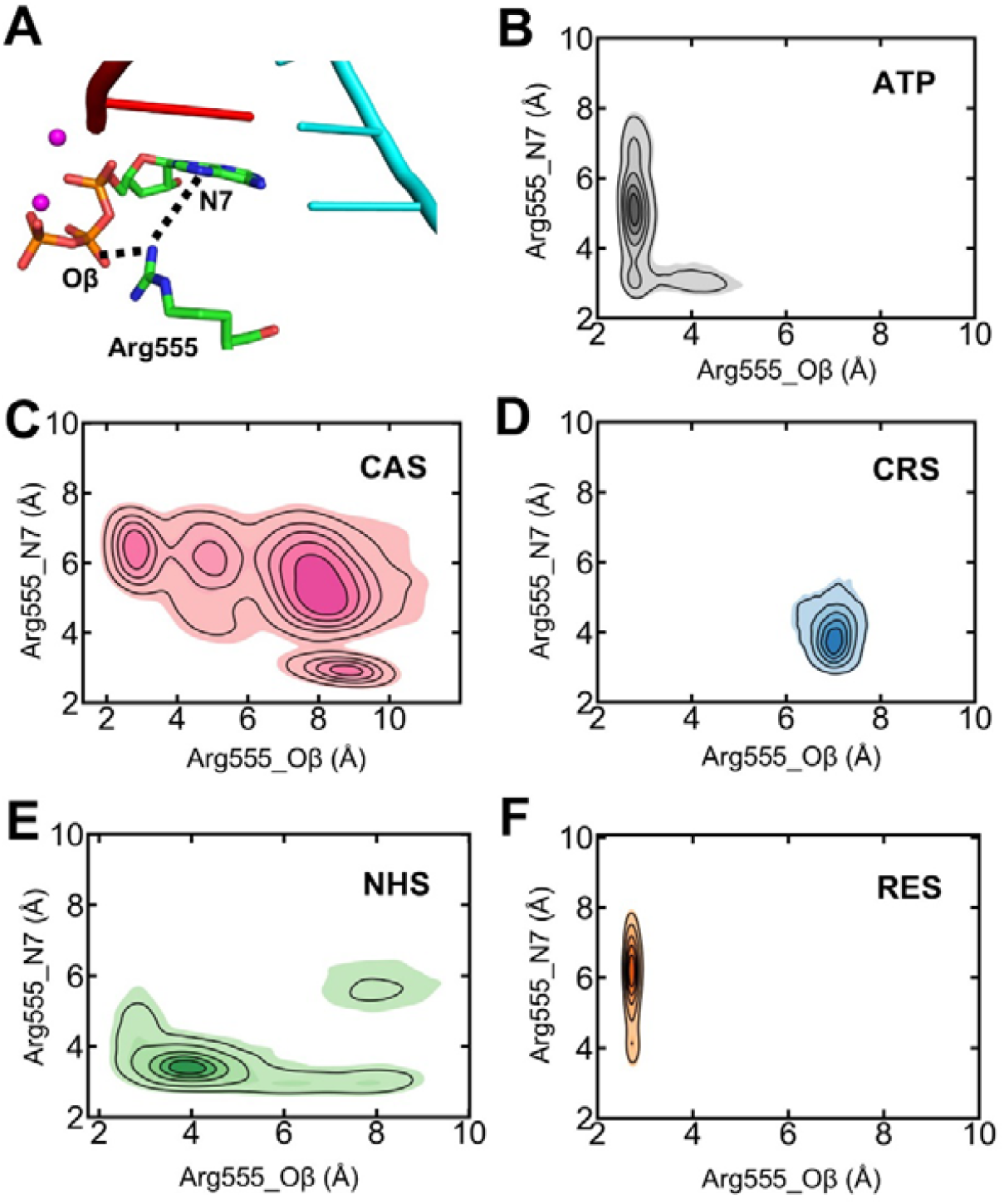
Investigation on the flexibility of Arg555 at the RdRp active site. (A) Diagram showing the distance between the side chain of Arg555 and the O_β_ and the nitrogen atom (N7) at the base ring. (B)-(F) Distance distribution of Arg555-O_β_ and Arg555-N7 of analog and ATP systems.

As a flexible residue at the RdRp active site, Arg555 responds diversely to the “thio” modification of different nucleotide analogs. It frequently interacts with O_Pβ_ in RES and ATP systems, while it is likely to interact with the base ring in the other three analog systems. This enhancement of Arg555-Base interaction caused by phosphate moieties torsion may affect the overall conformational changes of analogs at RdRp active site. The overall conformational change would directly impact the stability of the Watson–Crick base pairing. Meanwhile, the hydrogen bonds formed between chemical groups on the ribose and the surrounding residues are also affected. Previous studies revealed that a few conserved residues at the RdRp active site, including Asp623 and Asn691, may be critical in recognizing NTP binding at the RdRp active site^27, 28, 53^. Thus, we investigated the probability of hydrogen bonds formed by base pairing and ribose ring for analogs and natural ATP, shown in Figure 5. Five Hbonds were considered in the calculations, including two base pairing positions and three ribose positions (Figure 5A).

**Figure 5.**
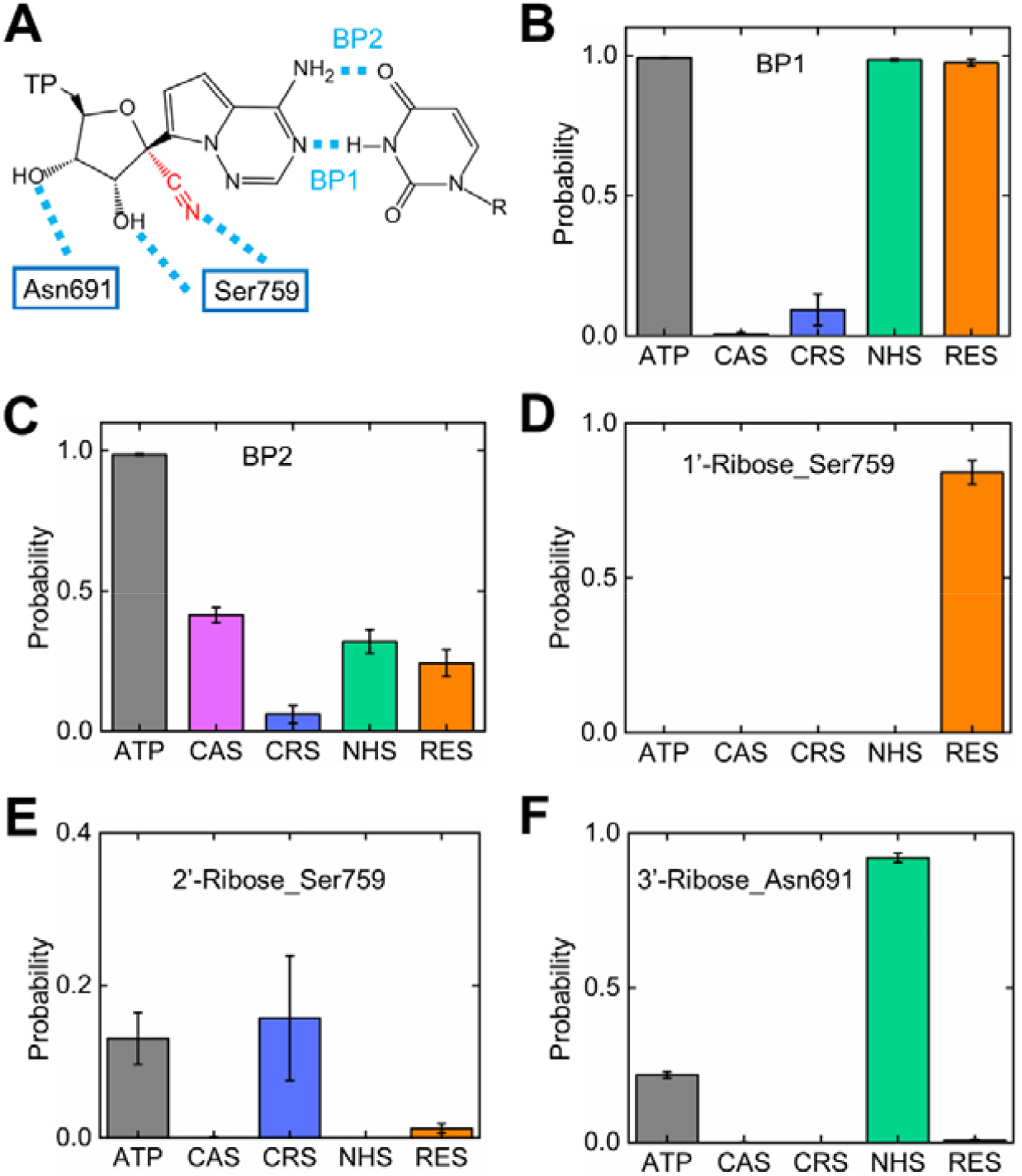
Investigation on the hydrogen bond (Hbond) formation between analogs and surrounding residues. (A) Diagram showing the base pairing of analogs and the Hbond defined by ribose and surrounding residues. (B) Hbond probability of base pairing of N…H-N. (C) Hbond probability of base pairing of N-H…O. (D)-(F) Probability of Hbond formed by ribose 1’, 2’ and 3’ positions and residue Ser759 and Asn691.

Natural ATP shows intact hydrogen bond interaction of base pairing at the RdRp active site. However, CAS and CRS barely have base pairings at the active site. In the CAS system, the hydrogen bond of N…H-N is not formed (Figure 5B), while the probability of N-H…O is 41.4±2.7% at the active site (Figure 5C). The probabilities of both hydrogen bonds in the CRS system are 9.3±5.6% and 6.0±3.1%, respectively. The first hydrogen bond (BP1) is kept intact for the NHS and RES systems, which are 98.4±0.3% and 97.4±1.2%, respectively. However, the second hydrogen bond (BP2) was reduced to 32.0±4.2% and 24.3±4.8%. Moreover, we checked the hydrogen bond formed by the ribose groups, including 1’, 2’, and 3’-ribose positions. We found that 3’-ribose of NHS formed a stable hydrogen bond with Asn691, and 1’-ribose of RES formed a stable hydrogen bond with Ser759. On the contrary, CAS and CRS formed very few hydrogen bonds with conserved residues at the active site.

The above hydrogen bond analysis demonstrated that NHS and RES formed more stable ribose Hbonds than CAS and CRS systems. This may explain their better structural stability at the RdRp active site. As discussed above, ribose modification of analogs would affect the binding positions of analogs at the RdRp active site through the surrounding residues. Previous studies provide diverse opinions on the role of these residues in stabilizing the natural NTP at the RdRp active site. Molecular dynamics research based on the SARS-CoV RdRp proposed that the hydrogen bond defined by the Asp623 and the 2’-hydroxyl groups contributed to the stability of NTP and analogs at the RdRp active site^28^, while Darst et al. argued that 2’-hydroxyl groups mainly interacted with Ser682 to maintain the position of GTP at the active site^37^. Moreover, a multiscale simulation study declared that the Ser682 forms a hydrogen bond with the nucleobase facing the incorporated GTP^65^. There is no apparent interaction between the ribose ring and surrounding conserved residues. Our study observed weak interaction at the ATP system between the 3’-hydroxyl group on the ribose and Asn691 with a probability of 21.8±1.1%. Meanwhile, Asp623 does not form a hydrogen bond with the 2’-hydroxyl group, which is consistent with a previous simulation study^65^. Thus, it is currently unclear at which stage the recognition process governed by the ribose hydrogen bond occurs, whether in the NTP incoming stage or the pre-catalytic stage. That is beyond the discussion scope of this paper, which requires more dynamic investigation or enhanced sampling to tackle the problem. Nonetheless, we confirmed that specific ribose modification offers structural stability for analogs in “thio” modification through hydrogen bonds with conserved residues at the RdRp active site.

Based on the above results, we found that the chemical groups on the 1’ and 3’-ribose positions play an important role in keeping appropriate positions of analogs in “thio” modification at the active site. In the NHS system, the probability of hydrogen bond defined by the 3’-hydroxyl group and Asn691 is much higher than in other analog systems, which provides more structural stability than CAS and CRS systems. The enhancement of the hydrogen bond also happened in the RES system, where interaction between the 1’-cyano group and Ser759 kept the analog stable at the active site. The 2’-ribose modifications, including the hydroxyl group on the CRS and the amino group on the NHS, do not form apparent interaction with surrounding residues, which play a minor role in the structural stability of analogs at the active site. Previous experimental and computational studies pointed out that analogs with 2’-ribose modification, such as Sofosbuvir, inhibit the nucleoside addition cycle of RdRp by disturbing the active site after being incorporated into the nascent strand instead of when it binds at the active site^10, 27, 31^. The methyl group on the 2’-ribose position of Sofosbuvir introduces steric hindrance for subsequent NTP and thus leads to immediate chain termination. Therefore, the strategy of 2’-ribose modification for further drug design is supposed to concentrate on the disruption of the RdRp active site when it is incorporated into the nascent strand instead of forming interaction with surrounding residues.

To further uncover the dynamical stability of analogs at the active site and the impact of analogs on the surrounding protein environment, we calculated the root mean squared fluctuation (RMSF) of each compound and natural ATP, as well as the motif of A-G. (Figure 7; Figure S2). Natural ATP shows an RMSF of 0.625±0.011Å at the active site, while CAS and CRS present higher RMSF of 0.783±0.008 Å and 0.814±0.051 Å, respectively. The NHS and RES present similar RMSFs, 0.592±0.006 Å and 0.599±0.020 Å, which are lower than the ATP and the other two analogs. Moreover, we checked the RMSF of different parts of analogs, including phosphate moieties, ribose ring, and base ring (Figure 6B-6D). We found that the RMSF of all three parts in the CAS and CRS is higher than that of other analogs and ATP, especially for the base ring of CRS. NHS and RES present lower RMSF on the ribose and base ring than other analogs and ATP. We also checked the RMSF of motif A-G of RdRp to investigate the impact of “thio” strategy of analogs on the protein environment. Motif F β hairpin loop was reported to stabilize the negative charges of phosphate moieties^66^. We found that motif F of analogs shows higher RMSF than natural ATP, except for the CRS system. Previous studies argued that motifs A and C are conserved in most viral RdRps^14^. For motif A, analog systems have higher RMSF than ATP except for the NHS, in which part of the residues show comparable RMSF to the ATP system. For motif C, analog systems show comparable RMSF to the ATP system (Figure 6E). We also calculated the RMSF of other motifs, as shown in Figure S2.

**Figure 6.**
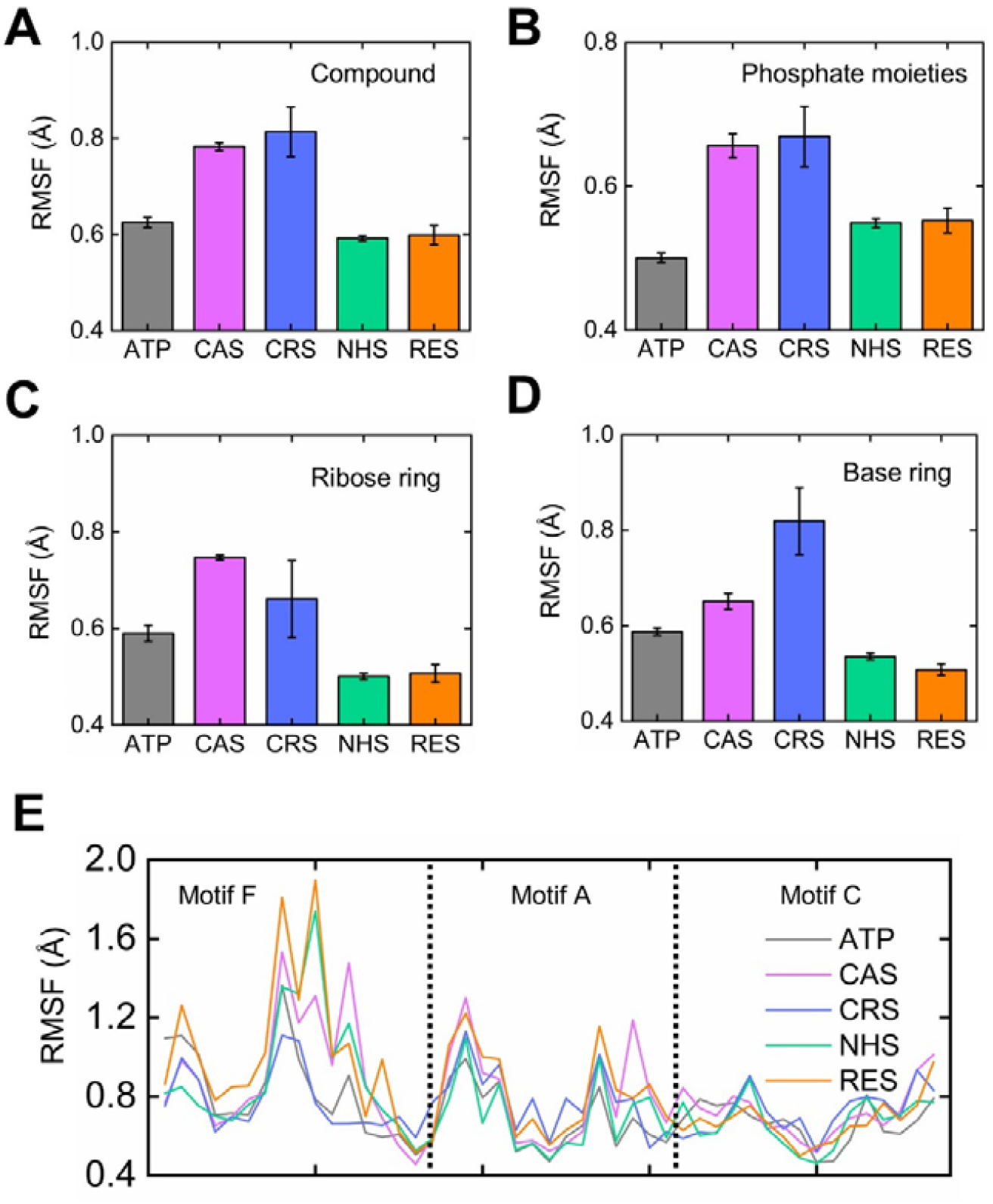
Investigation on the root mean square fluctuation (RMSF) on the analogs and surrounding conserved motifs. (A) RMSF of whole analogs compared with ATP. (B) RMSF of phosphate moieties. (C) RMSF of ribose ring. (D) RMSF of base ring. (E) RMSF of motifs F, A, and C.

**Figure 7.**
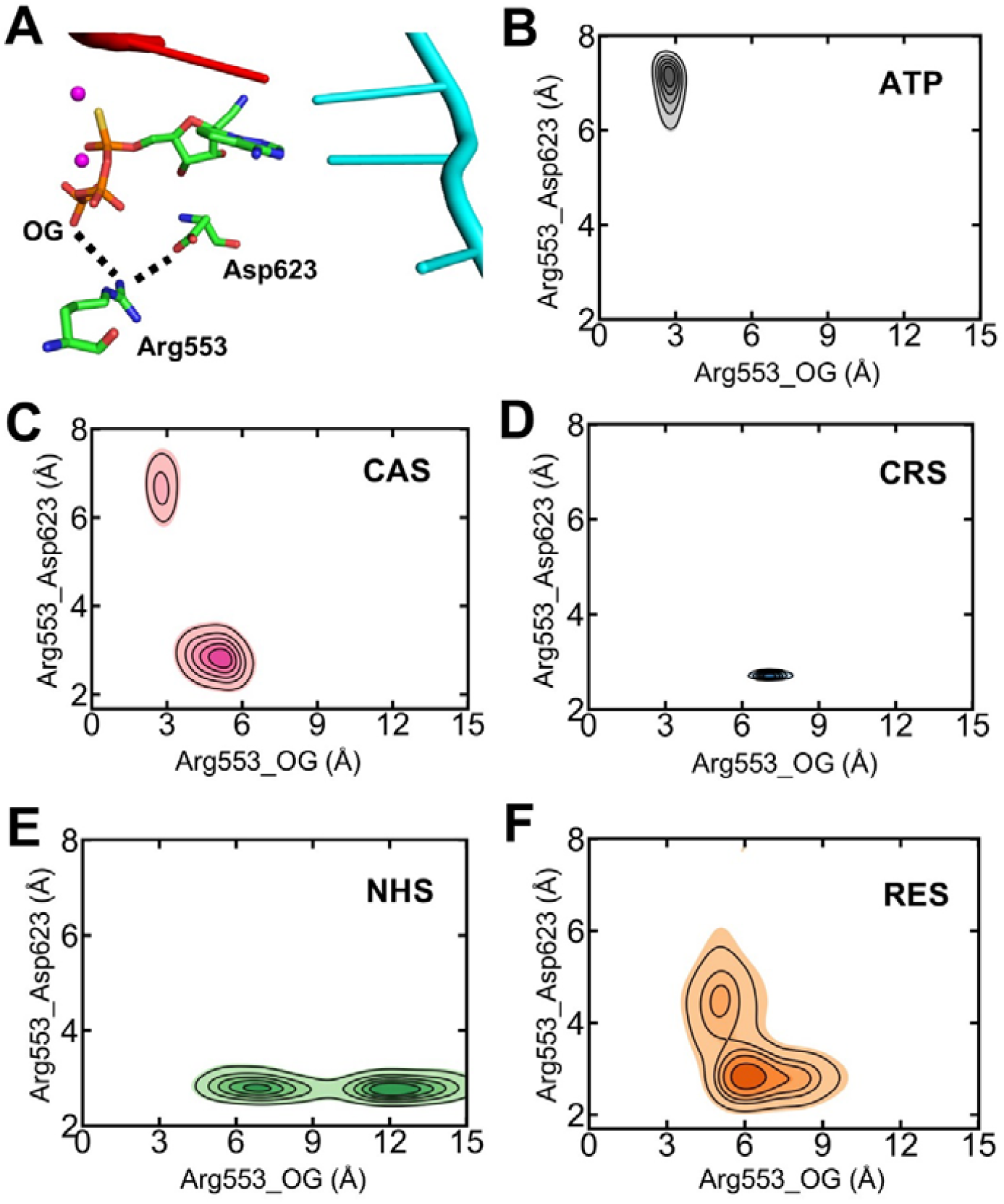
Investigation on the flexibility of Arg553 at the active site. (A) Diagram showing the RES, Arg553, and Asp623 at the active site. Side chains were selected to calculate the distance between residues. (B)-(F) Distance distribution of Arg553-O_G_ and Arg553-Asp623 of different analog systems and ATP system.

Residues on the motif F, such as Arg555 and Arg553, more frequently interacted with the phosphate moieties. We have shown that Arg555 is highly flexible to respond to the “thio” modification and can interact with the base of analogs. We further investigated how the torsion of phosphate moieties affects Arg553 on the motif F (Figure 7A). Arg553 tends to interact with Asp623 in the analog systems instead of interacting with the phosphate moieties (P_OG_). The distance of Arg553-P_OG_ in the natural ATP system is mainly concentrated at 2.5 Å (Figure 7B). In comparison, this distance is greater than 5 Å in analog systems, except for the CAS system, in which part of the conformations shares the same distribution with the ATP system (Figure 7C-7F). The distance of Arg553-Asp623 in the ATP system is greater than 6 Å in most conformations. This distance is less than 4 Å in most analog conformations. Thus, the conformational torsion caused by sulfur modification would profoundly affect the interaction mode between phosphate moieties and surrounding residues, leading to a higher RMSF of motif F.

Four analogs present diversity in the RMSF at the RdRp active site. “Thio” modification caused the apparent fluctuation in CAS and CRS systems, while it has limited impact on the NHS and RES. It is reasonable to propose that the lower RMSF of NHS and RES systems stem from the stable hydrogen bond interaction between ribose and surrounding residues. It should be noted that the 3’-hydroxyl group formed a stable hydrogen bond with Asn691 in the NHS system, while this interaction was not observed in the CAS system. We speculated that chlorine modification on the base ring of CAS corrupted the structural stability of analog due to its high electronegativity, which changed the overall position of CAS, reducing the probability of hydrogen bond between 3’-hydroxyl and Asn691.

To uncover the reason for the unfeasible stability of CAS and CRS, we further investigate the structural features and unexpected hydrogen bonds for the two systems compared with other analogs and ATP (Figure 8). In the CAS system, chlorine modification on the base ring destroyed base stacking at the *i* and *i*+1 sites on the template strand. The base ring at the *i*+1 site on the template strand is almost vertical to that of the *i* site. It also explains the probability of base pairing hydrogen bond of N…H-N at the active site reducing to zero in the CAS system (Figure 8B). We also checked the interaction between the CRS base ring and nucleoside at the *i*+1 site and observed an unexpected hydrogen bond on the nascent strand (Figure 8C). This unstable hydrogen bond with a probability of 41.6±6.3% disrupts the base stacking of the *i* and *i*+1 sites (Figure 8D). Meanwhile, the base pairing hydrogen bonds at the active site were ruined entirely due to the unfeasible base stacking, which also explains the higher RMSF of the CRS at the active site.

**Figure 8.**
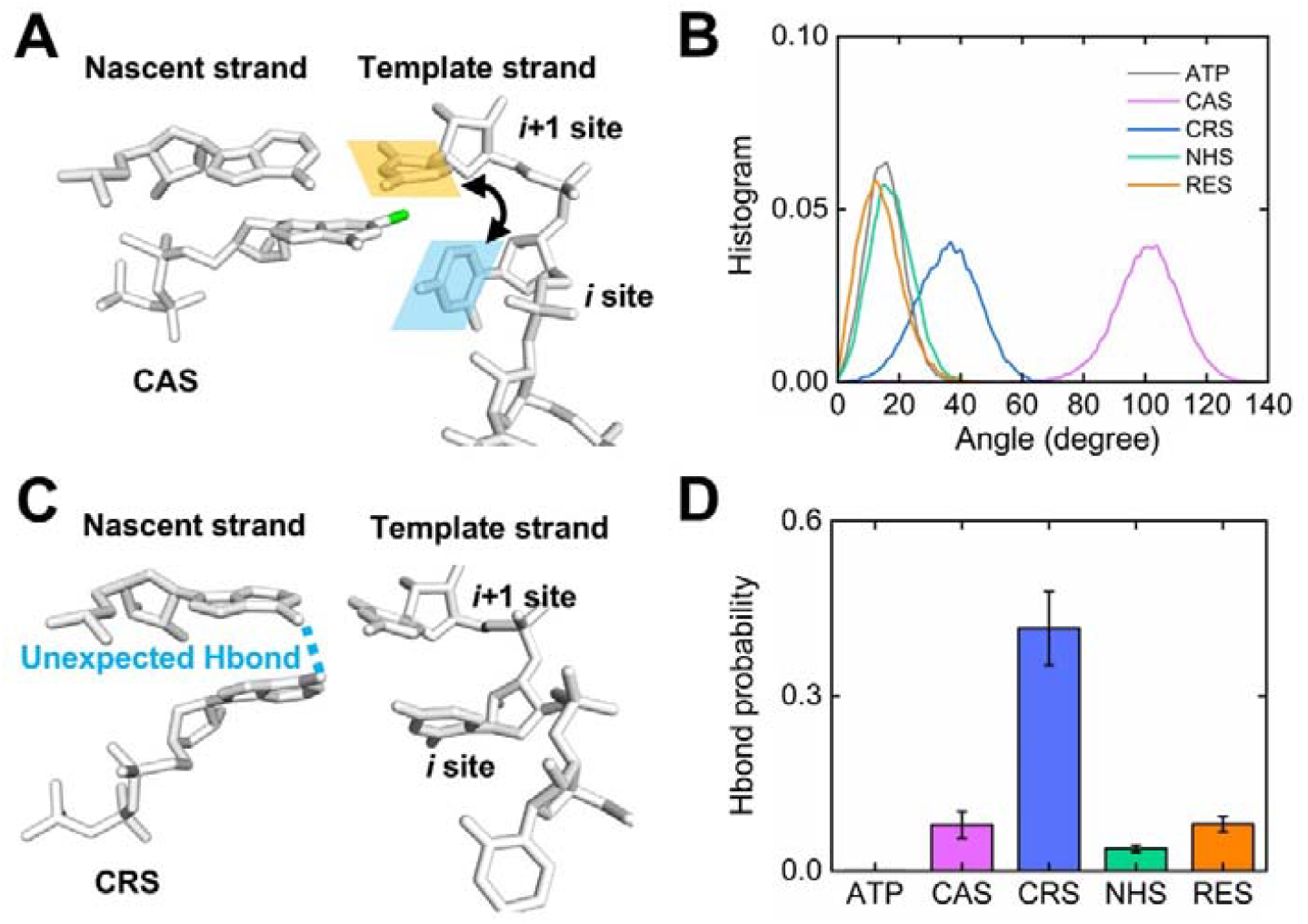
Investigation of the structural stability of analogs at the active site. (A) Diagram showing the CAS binding at the active site. The chlorine modification on the base ring is colored green. Two planes used to calculate the dihedral angle were defined by the base rings at the *i* site and *i*+1 site on the template strand. (B) Angle distribution of different analog systems compared with ATP system corresponding to (A). (C) Diagram showing the CRS binding at the active site. The unexpected hydrogen bond (Hbond) was colored blue. (D) Probability of unexpected Hbond of different analog systems and ATP system.

Therefore, the angle torsion of phosphate moieties caused by the sulfur atom modification can spread to the base ring conformations of analogs. With or without base modification, unexpected changes in base stacking would sabotage the catalytic conformation of analogs at the active site, which happened in CAS and CRS systems. On the contrary, hydrogen bond interactions between ribose and surrounding residues contribute to the structural stability of NHS and RES at the active site. We propose that polar groups on the ribose 1’ and 3’ positions are crucial to stabilizing the analogs at the active site with “thio” modification.

## Conclusion

In summary, we investigated the structural stability of four nucleotide analogs with sulfur modification on the phosphate moieties at the active site of SARS-CoV-2 RdRp to evaluate the feasibility of the “thio” strategy. Dynamical and structural analysis demonstrated that “thio” modification offers diverse structural stability for the analogs at the active site. The sulfur modification led to angle torsion on the phosphate moieties, which profoundly affected the analogs’ position at the active site and the interaction mode of surrounding residues.

Phosphate moiety torsion led to multiple structural changes of analogs at the active site. First, the distance of P_α_-O3 increased diversely in the analog system, reducing the catalytic ability of analogs and may lead to a lower reaction rate than the ATP system. This is consistent with previous experimental research that analog in “thio” modification decreased the catalytic rate compared to analog with triphosphate moieties^40^. Secondly, the base stacking at the active site changed unexpectedly. Consequently, the Watson-Crick pairing of analogs at the active site was broken to different extents. CAS and CRS systems presented inferior base pairing, especially for the CRS system. An unexpected hydrogen bond caused by structural change completely ruined the CRS system’s base pairing, increasing the RMSF of the analog at the active site. On the contrary, NHS and RES systems kept at least one hydrogen bond interaction of base pairing at the active site.

We propose that the hydrogen bond interaction between ribose and surrounding residues contributed to the structural stability of NHS and RES. The hydrogen bond between 1’-ribose and Ser759 stabilizes the RES at the active site, while the hydrogen bond between 3’-ribose Asn691 stabilizes the NHS. We emphasize the importance of hydrogen bond interaction related to the ribose in keeping feasible positions of nucleoside analogs with “thio” modification. No obvious evidence demonstrates that the 2’-ribose position plays a vital role in the structural stability of analogs in this study. Moreover, base modifications, especially the high electronegativity groups, should be chosen cautiously due to their potential to disrupt the base stacking.

Based on our data sample space, it is unlikely to propose a general rule to guide the ribose modification that benefits the structural stability of analogs at the active site. Nonetheless, we have shown that the molecular dynamics method revealed the interaction detail of analogs at the RdRp active site and emphasized the importance of ribose modification when introducing the “thio” strategy into the traditional nucleotide analogs. We confirmed that NHS and RES offer more feasible structural stability than CAS and CRS, with ATP as the reference. We suggest extensive MD simulations be performed on the potential analogs before further experimental research.

As a general strategy for modifying the nucleotide analogs, the “thio” modification offers a hopeful way to rescue the current analogs and avoid the excision from the SARS-CoV-2 nsp14. Our research revealed the dynamic and structural complexity of introducing multiple modifications into the analogs. Further experimental and theoretical studies should be performed to uncover the role of multiple chemical modifications on the performance of analogs at the active site of SARS-CoV-2 RdRp.

## Synopsis

Specific ribose modifications contribute to the structural stability of nucleotide analogs with α-thiotriphosphate at the active site of SARS-CoV-2 RdRp.

## Supporting information

Origianl_data

## Notes

The authors declare no competing financial interest.

## Acknowledgments

We acknowledge the financial support of this work by the Shandong Natural Science Foundation (No. ZR2024QB315).

## Data and Software Availability

Structure optimization in Hartree–Fock level for different analogs was carried out using the Gaussian 16 package. The AmberTools package was applied in the preparation of the parameters and input for MD simulations. All MD simulations were performed by the GROMACS 2022 package. Trajectory analysis was performed using the gmx module obtained in GROMACS, including gmx Hbond, gmx mindist and gmx gangle. The preparation of the structures and the images was carried out using PyMOL software, version 3.1.0. The complete workflow is reported in the “Methods” section of the manuscript. Original data and scripts, including input files, parameter files, topology files, and analysis scripts, were attached to the supporting information.

## Supporting Information

Distance distribution of two Mg^2+^ ions. Root mean square fluctuation (RMSF) of different motifs. Input files of MD simulations. Topology files and force field parameters of various analogs. Structures of complex composed of protein-RNA and analogs. Analysis scripts applied to calculate the HBond probabilities, RMSF, distance, and angles.

