## Supplementary material for "The Role of Ribose Modifications on the Structural Stability of Nucleotide Analogs with α-thiotriphosphate at the Active Site of SARS-CoV-2 RdRp": Origianl_data: all-comd.py

```
import numpy as np
fg=open('group.txt', 'r')
flines=fg.readlines()
for g in range(len(flines)):
    flines[g]=flines[g].strip()
fg.close()
namelist=np.loadtxt('name.txt',dtype=bytes).astype('str')
#namelist=['ATP']
index=np.loadtxt('index.txt', dtype=bytes).astype('str')
print(namelist)

for i in range(len(index)):
    for name in namelist:
        f = open('dis-command2', 'r')
        lines=f.readlines()
        atom=flines[i]
        lines[0]=lines[0].replace('chain B & r 235 & a NZ\\n13 & a O1A O2A\\nq\\n',atom)
        lines[0]=lines[0].replace('initial', index[i])
        lines[0]=lines[0].replace('SYS', name)
        lines[4]=lines[4].replace('SYS', name)
        lines[4]=lines[4].replace('initial', index[i])
#        if '3' in name:
#           lines[5]=lines[5].replace('24', '27')
#           lines[5]=lines[5].replace('25', '28') 
        ff = open('{}/{}/command'.format(index[i],name), 'w')
        ff.writelines(lines)
        f.close()
        ff.close()
```
