## Supplementary material for "The Role of Ribose Modifications on the Structural Stability of Nucleotide Analogs with α-thiotriphosphate at the Active Site of SARS-CoV-2 RdRp": Origianl_data: gene_just.py

```
import numpy as np
import matplotlib.cm as cm
from matplotlib.colors import ListedColormap
import seaborn as sns
import os
#from collections import OrderedDict
#import msmexplorer as msme
#import pickle
#import pandas as pd
#pd.set_option('display.max_row', None)
#import sys
#import heapq
from matplotlib import pyplot as plt
plt.switch_backend('agg')


#ind=int(sys.argv[1])
#namelist=['ATP','COP','CRP','FLP','VDP','DIP']
namelist=['VDP']
namelist2=['COP','CRP','FLP','VDP','DIP']
#xx=np.loadtxt('tmp1.txt',dtype=bytes).astype('str')
#yy=np.loadtxt('tmp2.txt',dtype=bytes).astype('str')
#xx_yy=['2prime_ENV', '3prime_ENV']

def boot_ind(micro):
    num=10
    ind=np.arange(1,num+1)
#    for k in xx_yy:
    for i in range(10):
#        if os.access("{}/total.dat".format(micro),os.F_OK)!=0:
        boot_random=np.random.choice(ind,num)
        np.savetxt('{}/rand_{}'.format(micro,i+1), boot_random, fmt='%d')
        data=[]
        for j in boot_random:
            rand=np.loadtxt('{}/results/{}-hb.dat'.format(micro,j), dtype='int')
            data.append(rand)
        data=np.asarray(data)
        data_con=np.concatenate(data)
        np.savetxt('{}/boot_{}-cut.dat'.format(micro,i+1),data_con,fmt='%d')

##############################
f_total=open('Total.dat','w')
for micro in namelist:
    boot_ind(micro)
#    for k in xx_yy:
    total_sum=[]
    for i in range(10):
        da=np.loadtxt('{}/boot_{}-cut.dat'.format(micro,i+1),dtype='int')
        pern=np.sum(da)/len(da)
        total_sum.append(pern)
    total_sum=np.asarray(total_sum)
    total_aver=np.sum(total_sum)/len(total_sum)
    total_std=np.std(total_sum)
    f=open('{}/zboot10just.dat'.format(micro),'w')
    f.write('{}\t{}'.format(total_aver, total_std))
    f.close()
    f_total.write('{}\t{}\t{}'.format(micro, total_aver, total_std))
f_total.close()
```
